# A combined analysis of genetically correlated traits identifies 107 loci associated with intelligence

**DOI:** 10.1101/160291

**Authors:** W. D. Hill, G. Davies, A. M. McIntosh, C. R. Gale, I. J. Deary

**Affiliations:** Centre for Cognitive Ageing and Cognitive Epidemiology, University of Edinburgh, Edinburgh, UK; Department of Psychology, University of Edinburgh, Edinburgh, UK; Division of Psychiatry, University of Edinburgh, UK; MRC Lifecourse Epidemiology Unit, University of Southampton, Southampton, UK

## Abstract

Intelligence, or general cognitive function, is phenotypically and genetically correlated with many traits, including many physical and mental health variables. Both education and household income are strongly genetically correlated with intelligence, at r_g_ =0.73 and r_g_ =0.70 respectively. This allowed us to utilize a novel approach, Multi-Trait Analysis of Genome-wide association studies (MTAG; Turley et al. 2017), to combine two large genome-wide association studies (GWASs) of education and household income to increase power in the largest GWAS on intelligence so far (Sniekers et al. 2017). This study had four goals: firstly, to facilitate the discovery of new genetic loci associated with intelligence; secondly, to add to our understanding of the biology of intelligence differences; thirdly, to examine whether combining genetically correlated traits in this way produces results consistent with the primary phenotype of intelligence; and, finally, to test how well this new meta-analytic data sample on intelligence predict phenotypic intelligence variance in an independent sample. We apply MTAG to three large GWAS: Sniekers et al (2017) on intelligence, Okbay et al. (2016) on Educational attainment, and Hill et al. (2016) on household income. By combining these three samples our functional sample size increased from 78 308 participants to 147 194. We found 107 independent loci associated with intelligence, implicating 233 genes, using both SNP-based and gene-based GWAS. We find evidence that neurogenesis may explain some of the biological differences in intelligence as well as genes expressed in the synapse and those involved in the regulation of the nervous system. We show that the results of our combined analysis demonstrate the same pattern of genetic correlations as a single measure/the simple measure of intelligence, providing support for the meta-analysis of these genetically-related phenotypes. We find that our MTAG meta-analysis of intelligence shows similar genetic correlations to 26 other phenotypes when compared with a GWAS consisting solely of cognitive tests. Finally, using an independent sample of 6 844 individuals we were able to predict 7% of intelligence using SNP data alone.

---

Intelligence, also called general cognitive function or simply *g*, describes the shared variance that exists between diverse measures of cognitive ability.^1^ In a population with a range of cognitive ability, intelligence accounts for around 40% of the variation between individuals in diverse cognitive tests’ scores.^2^ Intelligence is a heritable trait with twin- and family-based estimates of heritability indicating that between 50-80% of differences in intelligence.^3^ These genetic factors make a greater contribution to phenotypic differences as age increases from childhood to adulthood.^4^ Heritability estimates derived using molecular genetic data using the GREML-SC^5, 6^ method, indicate that around 20-30% of variation can be explained by variants in linkage disequilibrium with genotyped single nucleotide polymorphisms (SNPs).^7^ More recent methods such as GREML-KIN,^8^ and GREML-MS^9^, using imputed SNPs, have found that some of the heritability of intelligence can be found in variants that are in poor linkage disequilibrium (LD) with genotyped variants; by taking these into consideration, a heritability estimate of 0.54-0.50^10^ can be found. Intelligence is predictive of health states, including mortality;^11, 12^ a lower level of cognitive function in youth is associated with earlier death over the next several decades.^13^ Some of the association between intelligence and health is due to genetic variants that act across traits.^14, 15^

Few genetic variants have so far been reliably associated with intelligence differences.^16^ The sparsity of genome-wide significant single nucleotide polymorphisms (SNPs) discovered so far, combined with the heritability estimate, are indicative of a phenotype with a highly polygenic architecture, where the total effect of all associated variants is substantial, but each individual variant exerts a small influence. This is strong evidence that the lack of genome-wide significant loci associated with intelligence can be increased by increasing the sample size of GWASs as has been the case for phenotypes such as height,^17^ and schizophrenia.^18^

Two strategies have emerged in order to maximise power by increasing sample size for loci discovery in intelligence research. The first involves the meta-analysis of many GWAS conducted on intelligence.^19–21^ However, these GWASs are hampered by the fact that each individual sample tends to use different cognitive tests, and that the individual sample sizes are often small; therefore, even the resulting meta-analysis is underpowered to detect loci associated with intelligence. ^19, 21, 22^ This problem is ameliorated with studies like UK Biobank which contain a large number of individuals who have supplied genetic data as well as having taken cognitive tests^23^ that show high genetic correlations with intelligence^24^ (as derived from psychometrically validated test batteries).^16^

The second method is to use a “proxy” phenotype^25^ which is a phenotype that shows high phenotypic and genetic correlations with intelligence, and should therefore have a similar genetic architecture. Educational attainment has been successfully used as a proxy phenotype for intelligence^25^ owing in part to the ease at which it can be measured relatively consistently, facilitating the larger sample sizes required for loci discovery.^26^ Such methods have led to sample sizes of 293 723 for educational attainment, and the discovery of 74 loci attaining genome wide significance.^26^ The genetic correlation between the largest GWAS on intelligence and the largest GWAS on education is 0.70^16^.

In the present study we combine these approaches by using Multi-Trait Analysis of GWAS (MTAG)^27^ to meta-analyse summary statistics from genetically related traits. This enables us to add power to the GWAS on intelligence by adding in the genetic variance that is shared with proxy phenotypes to increase the sample size in the trait of interest here, i.e. intelligence. We use summary results from the largest available GWAS on intelligence (n =78 308).^16^ We combine it with the Social Science Genetic Association Consortium GWAS summary results on Education^26^ (n = 329 417, which include the individuals from UK Biobank). Finally we bring in the household income phenotype from UK Biobank^28^ (n = 96 900). Household Income shows a genetic correlation of r_g_ = 0.82 with education and r_g_ = 0.65 with the GWAS meta-analysis of Sniekers et al.^16^. These previous results show a high level of overlap of the genetic architecture of household income with education, and with intelligence satisfying the criteria for its use as a proxy phenotype.

By combining these three sets of GWAS summary results we increase the power to find loci associated with intelligence. The estimated effective sample size increased from 78 308 to 147 194. We then use bivariate linkage disequilibrium score (LDSC) regression^29^ to test whether these metaanalytic results have the same genetic architecture as other measures of intelligence. We use both SNP-based and gene-based GWAS to maximise our ability to discover loci and genes associated with intelligence before predicting phenotypic intelligence in an independent sample. Finally, we apply gene-set analysis using 10 891 gene sets sourced from Gene Ontology,^30^ Reactome,^31^ and, SigDB^32^ to derive biological meaning from our data. Our results indicate that, by drawing on multiple large GWAS data sets all measuring traits related to intelligence, we can attain greater statistical power to detect genetic variants associated with intelligence, facilitate our understanding of the underlying biology of intelligence differences, and make substantial phenotypic predictions of intelligence using SNP data.

## Method

### Samples

The data for the study were the summary statistics from three GWA studies conducted on cognitive and health/anthropometric traits. Supplementary Table 1 illustrates the samples used, and provides links to these data. Summary statistics were obtained from GWAS meta-analyses of intelligence (n=78 308),^16^ and education (n= 329 417),^26^ and a single large sample on household income (n = 96 900).^28^

In order to derive genetic correlations with intelligence and the proxy phenotypes (education, household income), as well as the final meta-analytic sample, we used summary statistics from 26 GWAS data sets. Supplementary Table 1 shows the data sets used and provides a reference and sample size for each data set used.

### Statistical analysis

#### Multi-Trait Analysis of GWAS (MTAG)

MTAG^27^ allows the meta-analysis of different traits that are genetically correlated with each other in order to increase power to detect loci in any one of the traits. Only summary data are required in order to carry our MTAG and, as bivariate LD score regression is carried out as part of an MTAG analysis to account for (possibly unknown) sample overlap between the GWAS results, these summary statistics need not come from independent samples. Our goal was to increase the power to detect loci associated with intelligence, and so the GWAS on intelligence by Sniekers et al^16^ was used as our primary GWAS data set. In order to add power to this data set we used two genetically correlated proxy phenotypes, i.e. Education^23^ (n = 329 417), and household income^25^ (n = 96 900). MTAG was run using the default settings.

### Clumping and annotation

Independent lead SNPs were identified using the clump function in PLINK. Index SNPs were defined as those that attained genome wide significance using an r^2^ threshold of 0.1 and a window size of 300kb. To map LD, the 1000 genomes phase 3 was used.^33^

### Cortical eQTL analysis

The web resource Braineac was used to examine evidence that the independent lead SNPs in each region had significant eQTL associations. Braineac uses data from UK Brain Expression Consortium (UKBEC) database containing post-mortem brains from 134 individuals who are free from any known neurological and neurodegenerative disorder.^34^ A total of 10 cortical regions were examined.

### Gene-based GWAS

Gene-based analysis was conducted using MAGMA.^35^ SNPs that were located within protein coding genes were used to derive a P-value describing the association found with intelligence. Gene locations and boundaries were used from the NCBI build 37 and LD was controlled for using the 1000 genomes phase 3 release.^33^ A Bonferroni correction was applied to control for the multiple tests performed on the 18 199 autosomal genes available for analysis.

### Tissue type gene expression

Differences in the expression rate genes identified in our SNP based and gene-based GWAS was conducted using FUMA.^36^ A total of 53 tissue types from the GTEx data base were examined using the average of the normalised expression per tissue by gene found in order to compare the level of expression of each gene across the 53 tissue types.

### Gene-set analysis

Gene-set analysis was conducted using MAGMA^35^ using competitive testing. A total of 10 891 gene-sets (sourced from Gene Ontology,^30^ Reactome,^31^ and, SigDB^32^) were examined for enrichment of intelligence. A Bonferroni correction was applied to control for the multiple tests performed on the 10 891 gene sets available for analysis.

### Genetic correlations

In order to test whether the genetic architecture of the meta-analysis of correlated traits conducted here produced a phenotype with the same genetic architecture as intelligence, we derived genetic correlations across 26 cognitive, socio-economic status (SES), mental health, metabolic, anthropometric, reproductive, and health and wellbeing phenotypes. We used linkage disequilibrium score regression^15^ to test whether each data set had sufficient evidence of a polygenic signal for a given phenotype. This was indicated by a heritability Z-score (Z_h_) of > 4 and a mean χ^2^ statistic of > 1.02. Following this, a minor allele frequency cut-off of < 0.01 was applied. Only SNPs that were in HapMap 3 with MAF > 0.05 in the 1000 Genomes EUR reference sample were included. Next, Indels and structural variants were removed along as were strand ambiguous variants. SNPs whose alleles did not match those in the 1000 Genomes were also removed. The presence of outliers can increase the standard error in LD score regression,^37^ and so SNPs where χ^2^ > 80 were removed. LD scores and weights for use with European populations were downloaded from (http://www.broadinstitute.org/~bulik/eur_ldscores/). A false discovery rate was applied to control for the number of tests performed.^38^ In the case of Alzheimer’s disease, a region encompassing 500 kb on each side of *APOE* was removed and the analysis re-run in order to ensure that the large effects in this region did not bias the regression.

### Genetic prediction

Genotype data were obtained from Generation Scotland: Scottish Family Health Study (GS:SFHS); (174) participants from GS:SFHS who also took part in UK Biobank were identified from their genetic data and removed from GS:SFHS. Following the removal of related individuals a total of 6 844 participants were available for prediction. A general factor of cognitive ability, i.e. a *g* factor, was derived using four cognitive tests: the Mill Hill Vocabulary Scale ((MHVS)^39, 40^, the Wechsler Digit Symbol Substitution Task (DST)^41^, Wechsler Logical Memory immediate and delayed added together^42^, and phonemic Verbal fluency (using letters C, F, L). ^43^ A general factor of intelligence was derived by extracting the first unrotated principal component from these four cognitive tests. This single component explained 42.3% of the variance and each of the individual tests used demonstrated strong loadings on the first unrotated component (DST 0.58, Verbal Fluency 0.72, MHVS 0.67, and Wechsler Logical Memory 0.63). We used scores on the first unrotated principal component as a measure of intelligence. The effects of age, sex, and population stratification (7 components^10^) were controlled for using residuals extracted from a regression model.

Using our meta-analytic data set on intelligence polygenic risk scores were derived for intelligence in GS:SFHS using PRSice.^44^ SNPs that were strand ambiguous and those with a MAF of p<0.01 were removed prior to deriving the polygenic risk scores. SNPs were clumped using the binary.ped files from the 6 844 participants in the GS:SFHS as a reference (r^2^ <0.1, 300kb window). Polygenic scores were then derived for each participant as the sum of alleles associated with intelligence, weighted by the effect size from our meta-analytic intelligence data set. A total of six polygenic risk scores were derived using the following P-value cut offs: 0.01, 0.05, 0.1, 0.5, and 1.

## Results

By meta-analysing the GWAS of intelligence^15^ with those of education^23^, and household income^25^ we were able to increase the mean χ^2^ in the intelligence data set from 1.197 to 1.370. This corresponds to an increase in the sample size from 78 308 to 147 194. The MTAG analysis that combined these three GWASs found 4 716 genome wide significant SNPs associated with intelligence (Figure 1). These SNPs were found in 107 independent loci, identified using the clump function in PLINK (Supplementary table 2),^45^ an increase of 89 loci compared to the Sniekers et al GWAS alone.^16^ By comparing the clumps found in the present study with the same SNPs in the intelligence, education, and household income data sets used in its construction we see low p-values across each of the three, consistent with the finding of a strong genetic correlation between each of the three phenotypes (Supplementary Table 3). Using the Braineac eQTL data base (http://www.braineac.org/), we found that of the 107 lead SNPs, 104 were found to have at least a nominal level of significance indicating their role as cis-eQTLs acting on cortical tissue (Supplementary Table 4) and confirming that our meta-analysis of intelligence, education, and income produces a phenotype with neurological underpinnings.

**Figure 1.**
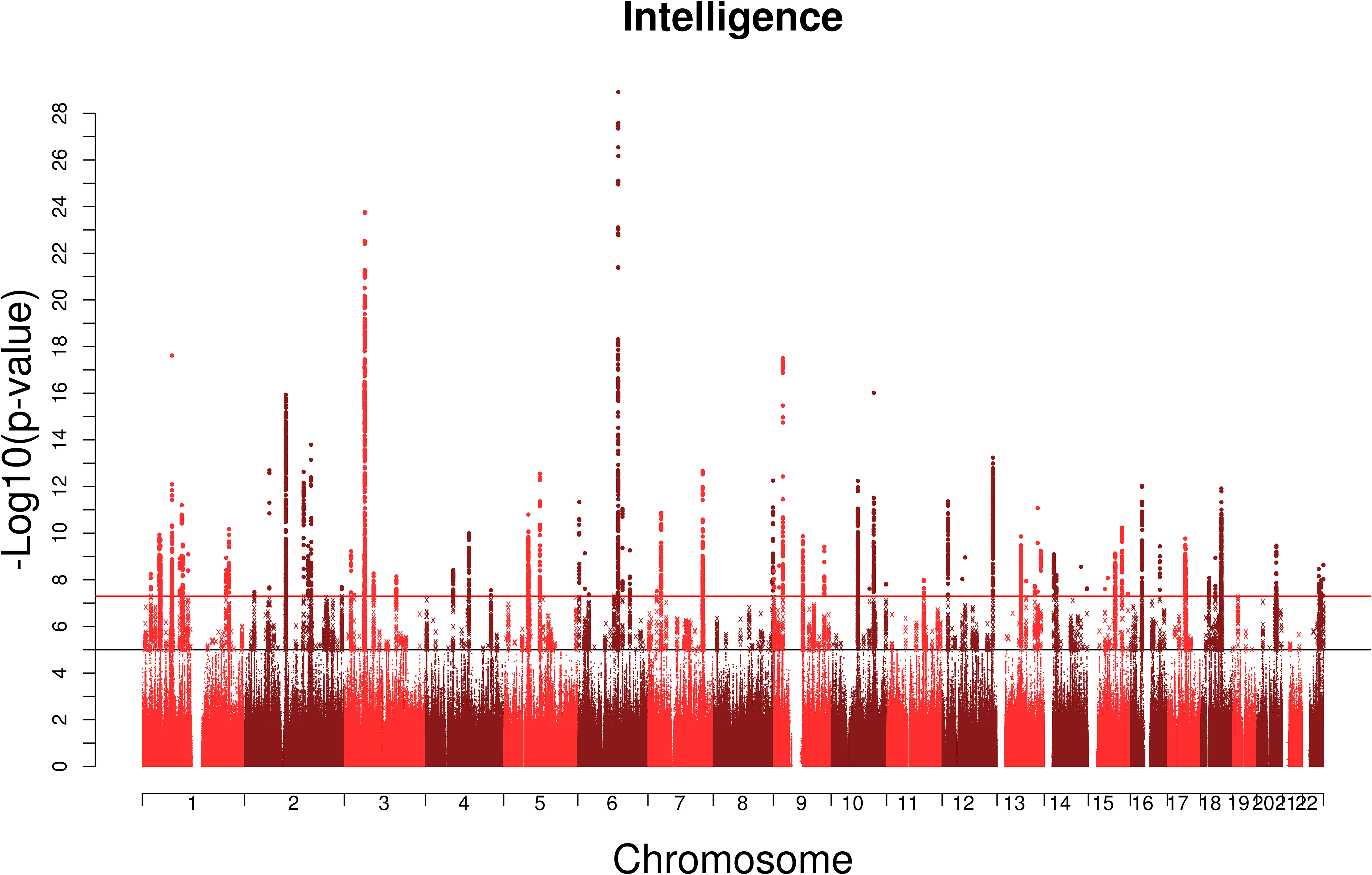
The results of our MTAG analysis. SNP-based GWAS Manhattan plot, negative log10 transformed P-values for each SNP are plotted against chromosomal location. The red line indicates genome-wide significance and the black line indicates suggestive associations.

A gene-based GWAS was conducted using MAGMA. Gene based analysis can increase power to detect significant associations as the signal across many SNPs (all within a gene) is combined.^46^ A total of 233 (Figure 2, Supplementary Table 5) genes attained genome-wide significance using gene-based GWAS. The genes identified using MAGMA were added to the genes implicated by the clumping analysis and after removing duplicate genes, a total of 338 genes were implicated as being involved in intelligence differences.

**Figure 2.**
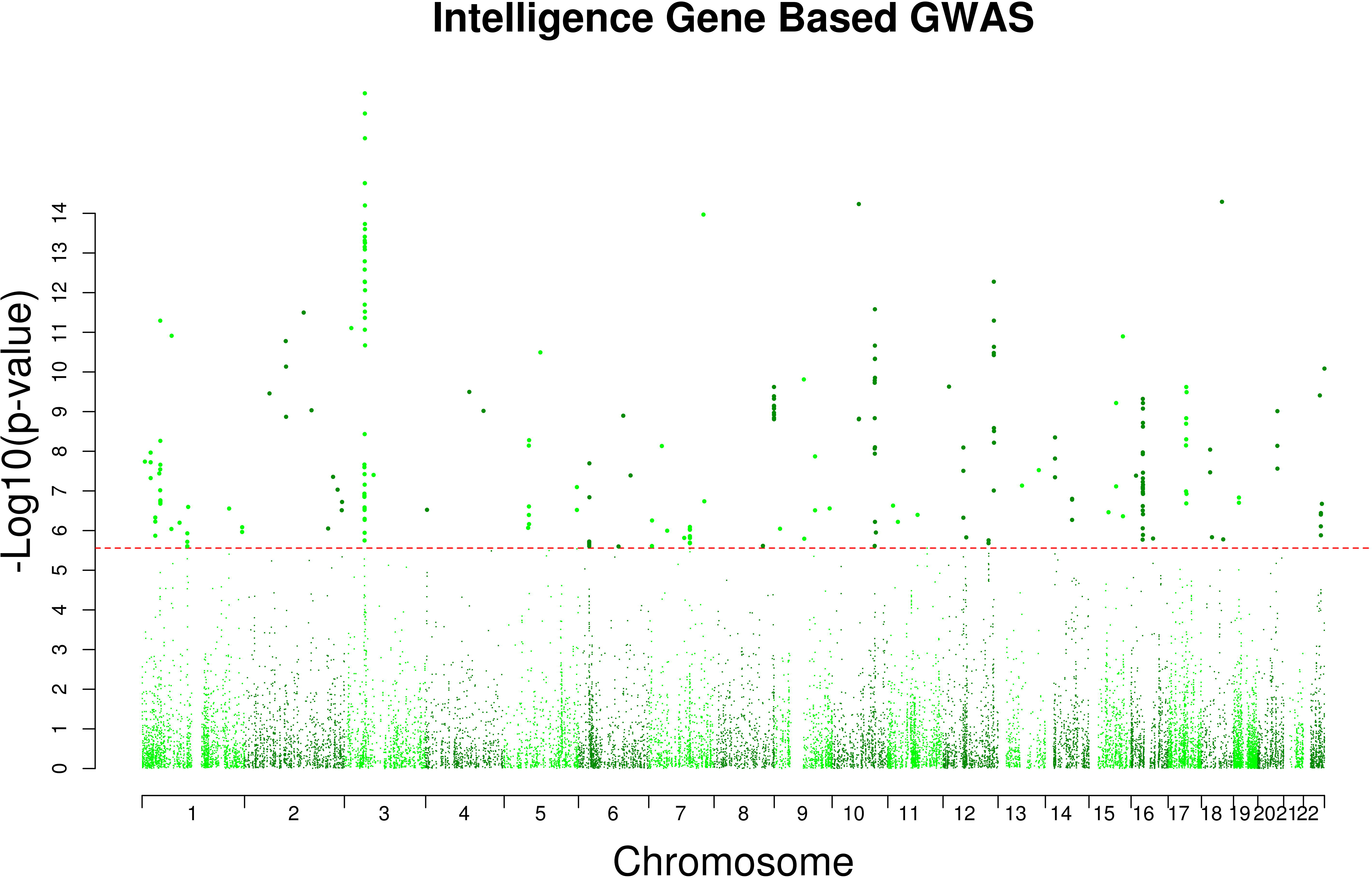
Gene based Manhattan plot, negative log10 transformed P-values for each gene (derived using MAGMA) are plotted against chromosomal location. The red line indicates genome wide significance.

Using our meta-analytic data set on intelligence we carried out polygenic prediction into GS:SFHS and found that 6.9% of phenotypic intelligence could be predicted (Table 2), an improvement of 43.75% on previous estimates of 4.8%.^16^ The polygenic risk scores that predicted the greatest amount of variance were those composed of the P <0.5, and P <1.0 cut offs indicating that despite our increase in power, many of the genetic variants associated with intelligence can still be found across the full distribution of P-values.

In order to obtain information on the biological systems involved in intelligence differences that are influenced by genetic variation, we conducted gene-set analysis using all genes available irrespective of their level of association. Using a competitive test of enrichment implemented in MAGMA, we identified three novel biological systems associated with intelligence differences and replicated previous findings by Sniekers et al^16^ (Table 1, Supplementary Table 6). Firstly, we identify a role for neurogenesis (gene-set size = 13 55 genes, P-value = 2.70×10^−7^), the process by which neurons are generated from neural stem cells. Secondly, a role was also found for genes expressed in the synapse (gene-set size =717 genes, P-value = 1.77×10^−6^), consistent with previous studies showing a role for synaptic plasticity.^47^ Thirdly, enrichment was found for the regulation of nervous system development (gene-set size = 722 genes, P-value = 1.88×10^−6^). Fourthly, the finding that regulation of cell development (gene-set size = 808 genes, P-value 9.71×10^−7^) was enriched for intelligence was replicated.^16^

**Table 1.**
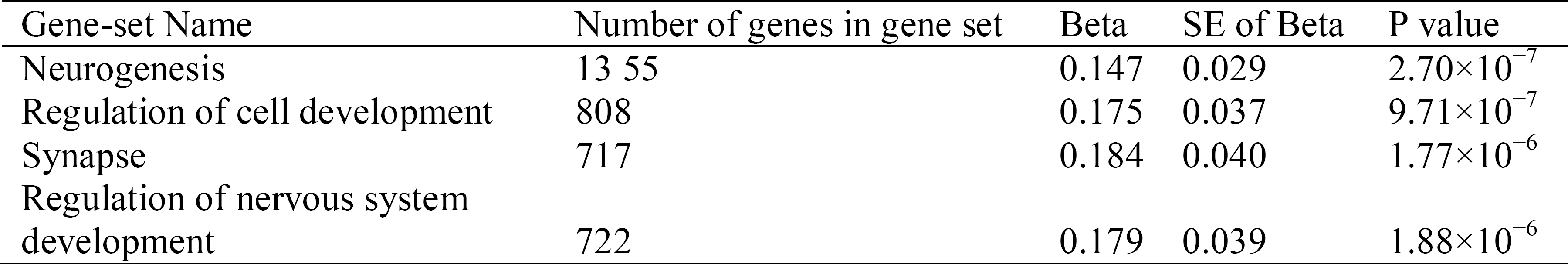
Gene-sets attaining statistical significance following Bonferroni control for multiple tests.

**Table 2.**
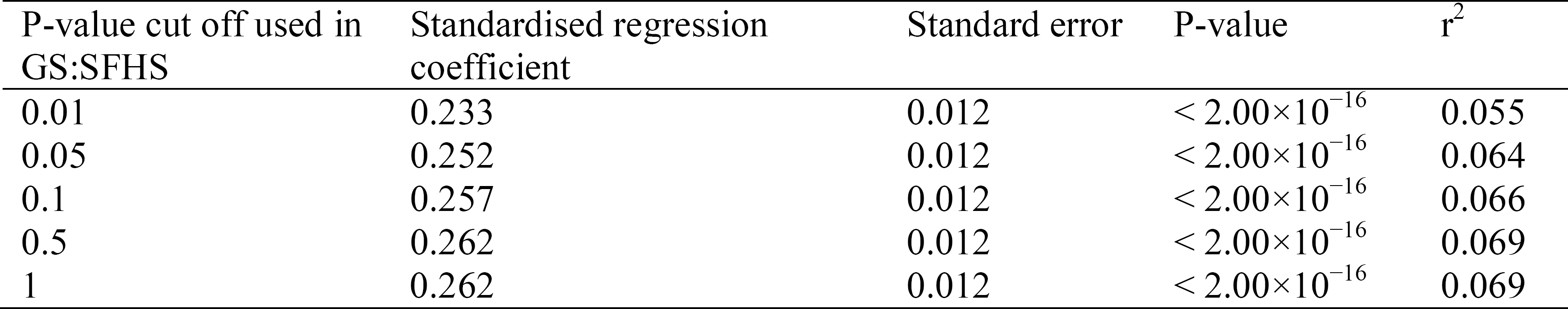
Showing the results of the polygenic risk scores analysis in GS:SFHS across each of the cut offs used.

The 233 genes identified by SNP based GWAS and gene based GWAS were investigated using 53 tissue types found in the GTEx data resource and implemented using Functional Mapping and annotation of genetic associations (FUMA).^36^ This tissue expression analysis indicated that over 100 of the genes showing association with intelligence are over-expressed in cortical tissues relative to the other tissues included in the FUMA analysis (Figure 3).

**Figure 3.**
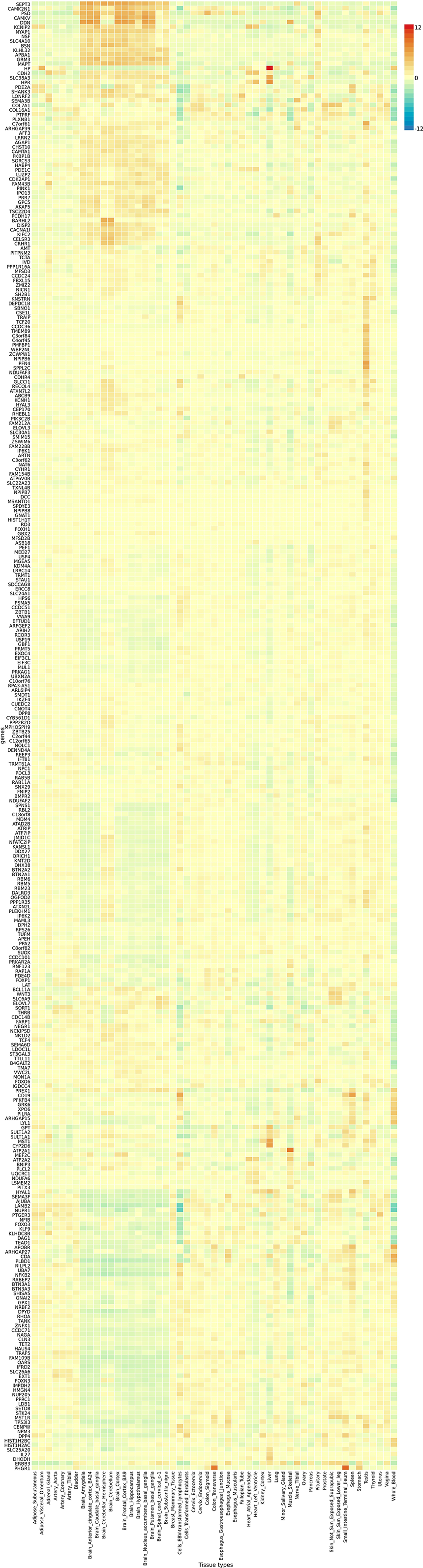
Heat map of gene expression for the 233 genes identified using clumping and using MAGMA across 53 tissue types. Values greater than zero (shown in red) indicate that a high level of expression relative to the mean level of expression across all tissues. Values less than zero (shown in blue) indicate that the mean level of expression was smaller than the mean across all tissue types.

In order to obtain evidence suggesting that the results of our meta-analysis produced a phenotype with the same genetic architecture as intelligence, we derived genetic correlations using 26 phenotypes (Figure 4, Supplementary Table 7). Many of these have been shown before using intelligence phenotypes^14, 29^, and replicated using the verbal-numerical reasoning phenotype from UK Biobank^48^, and some of these data sets are incorporated in the Sniekers^15^ meta-analysis; however, we include them to show the similarities and differences between the genetic architecture found in our meta-analytic intelligence data set and the three data sets used in its construction. These results are shown in Figure 4. We find a novel genetic correlation between intelligence and parental longevity; this is found using the intelligence^15^ GWAS (r_g_ = 0.33, SE = 0.08) and our meta-analytic sample (r_g_ = 0.44, SE = 0.07). This indicates that the polygenic load for greater intelligence is associated with greater longevity, using parental longevity as a proxy.

**Figure 4.**
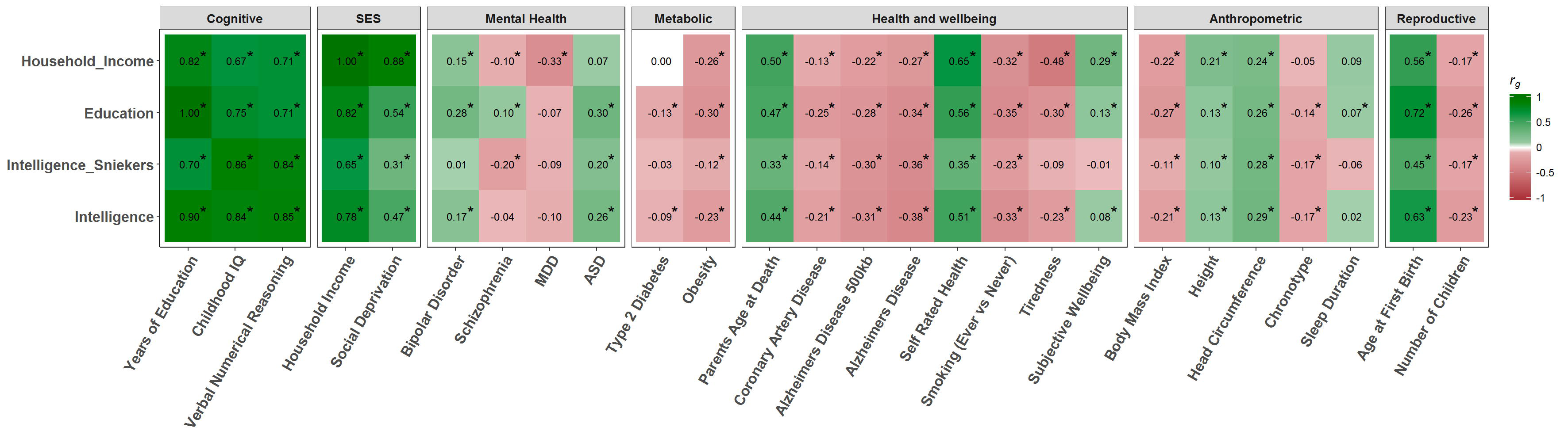
Heat map showing the genetic correlations between the meta-analytic intelligence phenotype, intelligence, education, and household income, with 26 cognitive, SES, mental health, metabolic, health and wellbeing, anthropometric, and reproductive traits. Positive genetic correlations are shown in green and negative genetic correlations are shown in red. Statistical significance following FDR correction is indicated by an asterisk.

When compared with cognitive phenotypes our meta-analytic data set showed strong positive genetic correlations with each of the three variables measured: childhood IQ, r_g_ = 0.84, SE = 0.06; years of education, r_g_ = 0.90, SE = 0.0005; and verbal numerical reasoning, r_g_ = 0.85, SE = 0.01. The point estimates for childhood IQ and verbal numerical reasoning are more similar for the intelligence GWAS^15^ (childhood IQ, r_g_ = 0.86, SE = 0.05, verbal numerical reasoning, r_g_ = 0.84, SE = 0.02) than for education (childhood IQ, r_g_ = 0.75, SE = 0.07, verbal numerical reasoning, r_g_ = 0.71, SE = 0.03). For the SES variables the point estimate for our meta-analytic intelligence data set fell between that of the intelligence^15^ GWAS and the Education GWAS^23^ which is a trend evident across all traits assessed.

For the mental health variables our meta-analytic intelligence data set showed a significant genetic correlation with bipolar disorder (r_g_ = 0.17, SE = 0.03) and with autism spectrum disorder (r_g_ = 0.26, SE = 0.04). The direction of these was both positive which, for autism spectrum disorder, was consistent with the results gained using established tests of intelligence.^14^ For bipolar disorder previous results have indicated a negative genetic correlation using established measures of intelligence, although after correcting for multiple tests this estimate was not statistically significant.^14^ Similar results were also found when examining schizophrenia, where no genetic correlation was found with our meta-analytic data set.

Differences between the GWAS on intelligence^15^ and our meta-analysis were also evident for tiredness, subjective wellbeing and type 2 diabetes. For these phenotypes, the genetic correlation is indistinguishable from zero for the intelligence^15^ GWAS but significant and in the same direction for both Education and our meta-analytic sample.

## Discussion

People with a higher level of cognitive function have been observed to have better physical and mental health, and to enjoy longer lives.^4, 11^ This paper exploited the high genetic correlations found between intelligence, education, and household income to add power to a GWAS on intelligence in order to find the loci, and biological mechanisms that help explain both intelligence, and the health differences with which it is associated. Through the use of summary statistics drawn from three large GWAS on intelligence and the proxy phenotypes of education and household income, and the recently developed method of MTAG^27^, we were able to assemble the sample sizes required to achieve the high levels of power needed to detect loci explaining differences in intelligence. These analyses produced a number of novel findings.

Firstly, we find 107 independent associations for intelligence in our GWAS, and highlight the role of 338 genes being involved in intelligence a substantial advance on the 18 loci previously reported.^16^ Of the 107 lead SNPs from these associations, 104 show significant signs of acting to produce expression differences in the brain and over 100 of these were over expressed in cortical tissue compared to other tissue types.

Secondly, we use our meta-analytic GWAS data to predict almost 7% of the intelligence differences in an independent sample. Previous estimates of prediction have been around 5%^16^ indicating that prediction accuracy can be improved by drawing on existing data sets of proxy phenotypes for intelligence. Additionally, polygenic scores derived using the MTAG method can be used to make meaningful predictions regarding an individual level of intelligence.

Thirdly, we report the novel finding that the polygenic signal across our GWAS data set clusters in genes involved in the process of neurogenesis, genes expressed in the synapse, and genes involved in the development of the nervous system, providing a theoretical rationale as to how genetic differences can result in the physiological differences that are part of the biology of intelligence differences.

The finding of neurogenesis gene-set enrichment for intelligence is a persuasive finding as neurogenesis has been linked to cognitive processes in rodent model particularly pattern separation and cognitive flexibility. New neurons are continually made in humans in the subgranular zone of the hippocampus and in the striatum.^49^ In rodent studies, experimentally reducing neurogenesis results in a decrease in the ability to discriminate between highly similar patterns,^50^ whereas by increasing the number of new neurons produced results in an increased ability to successfully discriminate between highly similar stimuli.^51^ Additionally, neurogenesis appears to be involved in cognitive flexibility by serving to avoid interference between novel and previously formed memories in a spatial navigation task.^52, 53^ Such findings have been expanded to include touch-screen discrimination tasks,^54^ as well as active place avoidance.^55^ Across these experiments the common finding was that neurogenesis was not required for the learning of the task but, rather, for the reversal of the rule once the formally correct response had changed, suggesting that neurogenesis is an important mechanism for cognitive flexibility. Replication of this finding for enrichment of neurogenesis in intelligence GWAS data in an independent sample is required to confirm this finding of a biological mechanism associated with intelligence differences in humans.

Finally we show using genetic correlations with 26 other traits that our meta-analytic intelligence GWAS has a highly similar genetic architecture to intelligence alone. Where our phenotype did differ was where education and intelligence, and household income differ. This was most evident for traits of bipolar disorder and schizophrenia where positive genetic correlations have been observed for education but negative for intelligence.^14, 29^ Our new findings provides evidence that the differences in genetic correlations between intelligence and education found for traits such as schizophrenia^14^’ ^48^, and bipolar disorder, are due to genetic effects that act solely on intelligence being those that are negatively genetically correlated with bipolar disorder and schizophrenia indicating their protective against these disorders. However, the genetic variants that act across each of these traits are those that show positive genetic correlations with both. By meta-analysing intelligence, with the genetic component of education that overlaps with intelligence, the relative contribution of variance that is unique to intelligence lessens, and so too does the magnitude of these genetic correlations.

This limitation, a greater proportion of variance that is common across education and intelligence in our results, has implications for the results of our GWAS, as those variants that benefit the most from meta-analysis across genetically correlated traits will, by definition, show association with each trait in our meta-analysis. Whereas the final results of our GWAS do indicate the loci that are involved in intelligence differences, our GWAS will be overrepresented by effects that are also associated with education and household income. Nevertheless we did find associations on chromosomes 15, 16, 18, and 20 that were not found in the latest GWAS on education,^56^ and peaks that were identified as being associated with education, on chromosomes 19, and 21^56^ were not found to be genome-wide significant in our meta-analytic data set. Future work using large GWAS that are exclusively based on established tests of intelligence^21^ will provide a valuable sample in which to replicate and validate these findings. The strength of the MTAG approach used here, drawing additional power from related phenotypes, lies in the accumulation of additional power to detect loci, make more accurate predictions based on SNP data, and the ability to identify the biological significance behind the polygenic signal in such data sets.

## Acknowledgements

This work was undertaken in The University of Edinburgh Centre for Cognitive Ageing and Cognitive Epidemiology, supported by funding from the Biotechnology and Biological Sciences Research Council (BBSRC) and the Medical Research Council (MRC; MR/K026992/1) and the University of Edinburgh. This funding supports IJD, GD and CRG. AMM and IJD acknowledge the support of the Wellcome Trust (Wellcome Trust Strategic Award “STratifying Resilience and Depression Longitudinally” (STRADL) Reference 104036/Z/14/Z).

This research has been conducted using the UK Biobank Resource under UK Biobank application 10279.

WDH is supported by a grant from Age UK (Disconnected Mind Project).

The authors declare no biomedical financial interests or potential conflicts of interest.

